# Dependable Algorithm for Visualizing Snoring Duration through Acoustic Analysis

**DOI:** 10.1101/2021.05.18.444630

**Authors:** Hsueh-Hsin Kao, Yen-Chang Lin, Yee-Hsin Kao, Madan Ho, Hsiao-Chen Yu, Chun-Lung Wang, Jui-Kun Chiang

**Author notes:** **Co-correspondant author:** Dr. Jui-Kun Chiang, Department of Family Medicine, Dalin Tzu Chi Hospital, Buddhist Tzu Chi Medical, Foundation, 2, Minsheng Road, Dalin 62247, Chiayi, Taiwan. **Funding information** JK Chiang received grants from the Dalin Tzu Chi Hospital, Buddhist Tzu Chi Medical Foundation (DTCRD108(2)-E-09).

## Abstract

**Background:** Snoring is a nuisance for the bed partners of people who snore and is also associated with chronic diseases. Estimating the snoring duration from a whole-night-sleep period is challenging. The authors present a dependable algorithm for visualizing snoring durations through acoustic analysis.

**Method:** Both instruments (Sony digital recorder and smartphone’s SnoreClock app) were placed within 30 cm from the examinee’s head during the sleep period. Subsequently, spectrograms were plotted based on audio files recorded from Sony recorders. The authors developed an algorithm to validate snoring durations through visualization of typical snoring segments.

**Results:** In total, 37 snoring recordings obtained from six individuals were analyzed. The mean age of the participants was 44.6 ± 9.9 years. A 3-s segment demonstrated the typical dominant frequency bands and amplitude waves of two snores. Every recorded file was tailored to a regular 600-s segment and plotted. Visualization revealed that the typical features of the clustered snores in the amplitude domains were near-isometric spikes (most had an ascending–descending trend). The recorded snores exhibited one or more visibly fixed frequency bands. Intervals were noted between the snoring clusters and were incorporated into the whole-night snoring calculation. The correlative coefficients of snoring rates of digitally recorded files examined by Examiners A and B were higher (0.865, *p* < 0.001) than those with SnoreClock app (0.757, *p* < 0.001; 0.787, *p* < 0.001, respectively).

**Conclusion:** A dependable algorithm with high reproducibility was developed for visualizing snoring durations.

## 1. Introduction

Snoring, defined as snorting or grunting sounds during sleep, is caused by the vibration of soft tissues throughout the upper airway. Snoring is not only a nuisance for the bed partners but also associated with chronic diseases.[1] A study reported that the prevalence rate of chronic snoring is higher in adult men (40%) and lower in adult women (20%), although the variation is large.[2] The snoring prevalence is more than 40% in Asian countries, including Taiwan (59.1%),[3] Malaysia (47.3%),[4] and Turkey (40.7%).[5] One of the reasons for varying snoring prevalence might be that people of Chinese descent tend to have narrower cranial bases and flatter midface structures than do people of other races.[6]

Studies have proposed the criteria that all snoring sounds measured through manual audio recording should have an audible oscillatory component and be synchronous with breathing and yet protuberant from background sounds. [7, 8] Generally, sounds have two characteristics: sound tone (frequency) and sound intensity (amplitude). Similarly, snoring can be defined through sound intensity and frequency. A study defined snoring as a breathing sound intensity of >25 dB.[9] Furthermore, typical breathing sound frequencies for snoring range from 110 to 190 Hz, although frequencies even from 800 to 5000 Hz have been reported.[10-13] Therefore, some researchers have transferred collected audio data to Mel-frequency cepstral coefficients and have used hidden Markov models to detect and monitor snoring by using audio data. The reported detection accuracy of snoring from using an ambient microphone range from 87% to 98%. [9, 10, 14, 15] However, the aforementioned studies were performed in sleep laboratories.

Subsequently, a study suggested a snoring epoch method for estimating snoring duration according to a threshold level of the maximum amplitude for every 30-s segment. A 30-s sleep epoch that contained three or more snoring signals was deemed a snoring epoch. The snoring duration was subsequently calculated as the sum of the snoring epoch * 30 s. A shortcoming of this epoch method is that different threshold levels may output different results.[16] Another method for determining snoring duration is manual listening. However, this method might yield a large variation in results due to the difficulty of concentrating on hearing over a long period.

Snore apps are software applications that run on smartphones and record sound information while the user is sleeping, and they have provided convenient and personalized sleep care.[17] With the progression of innovative monitoring mechanics and techniques, the apps can be used at home and examinations can be performed as often as every night without interruptions to the user’s sleep. Studies have reported that the precision of smartphone apps for predicting snoring ranges from 93% to 96%, although apps can vary greatly among smartphone models.[18] SnoreClock was one of smartphone applications and it has a high predictive value for snoring.[19]

In this study, the authors present a dependable algorithm for calculating snoring duration on the basis of the typical features of snoring frequency and amplitude and enable the visualization of snoring duration on a spectrogram. The authors compared the results of using our trusted method with those of using the home-based SnoreClock app.

## 2. Materials and Methods

### 2.1 Study participants

Six individuals with habitual snoring voluntarily participated in this study. The study was conducted over different periods at their respective homes in Taiwan. Informed consent was obtained from all participants in this study. The study protocol was reviewed and approved by the Research Ethics Committee of the Buddhist Dalin Tzu Chi Hospital in Taiwan (No. B10703013).

All participants placed their smartphones and digital recorders (Sony ICD-SX 2000, Sony Electronics Inc., Tokyo, Japan) away from them yet within arm’s reach before going to sleep. The smartphones and recorders were placed at the cranial side of the shoulder within 30 cm from the head to optimally record the snoring sounds of patients. Snoring rates were then determined through measurement of snoring durations over the whole sleep period. In this study, the snoring durations were composed of snoring signals and interval pauses of no more than 100 s.

### 2.2 Snoring rates obtained using our dependable algorithm

The dependable algorithm was obtained through analysis of files obtained from Sony recorders. Snoring sounds throughout the night were recorded using portable digital sound recorders with linear pulse-code modulation (ICD-SX 2000, Sony Electronics Inc., Tokyo, Japan). The manufacturer of the sound recorder was not involved in this study. This recorder has two built-in high-performance electric dynamic microphones. These microphones were moved to a 90-degree “X-Y pattern.” The low-cut filter and limiter switches were set to the “OFF” position. The acquired audio signals were then digitalized at a sampling frequency of 44.1 kHz, PCM (pulse-code modulation), and 16 bits per sample. The authors subsequently moved these files to our computer for further evaluation. The files were records taken every 600 s. Dominant frequencies for each 0.01-s segment were analyzed and plotted. Simultaneously, Hilbert amplitude envelopes were smoothed using a sample-moving average of 1000 samples with a 25% overlapping sliding window. [9, 10, 20, 21]

The R codes (R Foundation for Statistical Computing, Vienna, Austria) used are presented as follows.

To read the audio file (190402_0310.wav in E disc, for example):

>library(tuneR)

>file.1 <-readWave(“E:/190402_0310.wav”)

To cut the file into 600-s segments:

>file.1.600sec <-cutw(file.1, from =0, to=600, output=“Wave”)

The first part of the 600-s segment was, for example:

For the frequency domain:

>df.plot <-dfreq(file.600sec.1, at = seq(1, 599, by = 0.01))

For the amplitude domain:

>amp.plot <-timer(file.600sec.1, threshold=70, msmooth=c(1000,25), dmin=0.001,

envt=“hil”, plot=TRUE)

Visualization revealed that the clustered snores in the amplitude domains generally exhibited near-isometric spikes with a trend of initial ascension and subsequent descension. Moreover, the frequency domains of the snoring records exhibited one or more visibly fixed frequency bands. Thus, the authors combined the correspondent two figures (matched amplitude domains and frequency domains) as snoring clusters. Notably, intervals (no more than 100 s) lay between the snoring clusters, and they were both incorporated into the whole-night snoring duration. Plots with typical amplitude domains and frequency domains classified as snoring that did not meet our criteria were classified as “noises”. For the aforementioned reasons, the authors set these guidelines, which enabled visualization of snoring durations. To test the reproducibility of the method of calculating snoring duration by using Sony digital recorders, two researches thoroughly examined the digitally recorded files.

The “noise” patterns did not possess the aforementioned typical features of snoring sounds. Therefore, the authors could easily differentiate noises (e.g., those from a fan, an air-conditioner, a toilet, outdoors, coughing, and groaning) from genuine snoring sounds. In the case that snoring durations were ambiguous, the authors carefully listened to the recorded files for confirmation.

### 2.3 Determining snoring rates by using smartphone apps

Because smartphone apps are accessible and portable, the SnoreClock app for smartphones was used. The snoring rates were displayed on the screens of smartphones. The developers of SnoreClock app were not involved in this study.

The authors could visualize typical snoring durations according to the typical frequency and amplitude of snoring on spectrograms. All snoring durations were summed, and snoring rates were determined.

### 2.4 Statistical analysis

The R statistical software, version 3.4.1, was used for all statistical analyses. The seewave package for R was applied. Hilbert amplitude envelopes were smoothed using a sample movement average of 1000 samples, with a 25% overlapping sliding window based on the timers’ function. Dominant frequencies were processed every 0.01 s through fast Fourier transform using the dfreq function of the R package.

Statistical significance was set at *p* < 0.05, and all tests were two-tailed. Continuous variables were presented as mean ± standard deviation where t-test or Wilcoxon test were indicated by cases. Correlation analysis was performed to evaluate associations in snoring rates between methodologies and examiners. ANOVA was used to test the differences between three groups of continuous variables.

## 3. Results

In total, 37 recordings from six participants (two female and four male) were used for the analysis. Their mean age was 44.6 ± 9.9 years. The mean recording time was 5.73 ± 1.18 h. (Table 1) The sound files captured with a digital recorder were transferred to a computer for analysis. The depicted bands of typical snoring frequency could vary during the recording period at the intraindividual and interindividual level. A 3-s period demonstrated the dominant frequency bands and amplitude spikes of the typical snoring duration. The three mean frequency bands of snoring presented on spectrograms were 4694, 2713, and 1083 Hz. The fourth mean frequency band of snoring was hidden in background noise (Figure 1). Every recorded file was segmented at 600-s (10 min) intervals. The dominant frequencies for every 0.01 s were calculated through fast Fourier transformation and plotted, and the Hilbert amplitude envelopes were plotted as well. Under this time scale, the matched 10-min plots of frequency bands and amplitude spikes were combined to estimate snoring durations. Isometric spikes (amplitude domain) that formed during participants’ snoring and that were as long as the fixed frequency bands (frequency domain) were paired and then grouped into isolated clusters, generally at intervals (typically less than 100 s). Therefore, the repeated “cluster–interval–cluster–interval” pattern must be identified to calculate snoring duration while “noises” are simultaneously excluded. The amplitudes could vary with time. Figure 2 shows an example of the typical duration of a 600-s (10 min) snore. The entire snoring duration was composed of summed interspersed signals and intervals (typically less than 100 s). Noises (such as those from walking, shifting position in bed, doors opening or closing, waterflow, talking, and curtains opening or closing) did not have typically featured fixed frequency bands and repeated amplitudes (Figure 3).

**Table 1.**
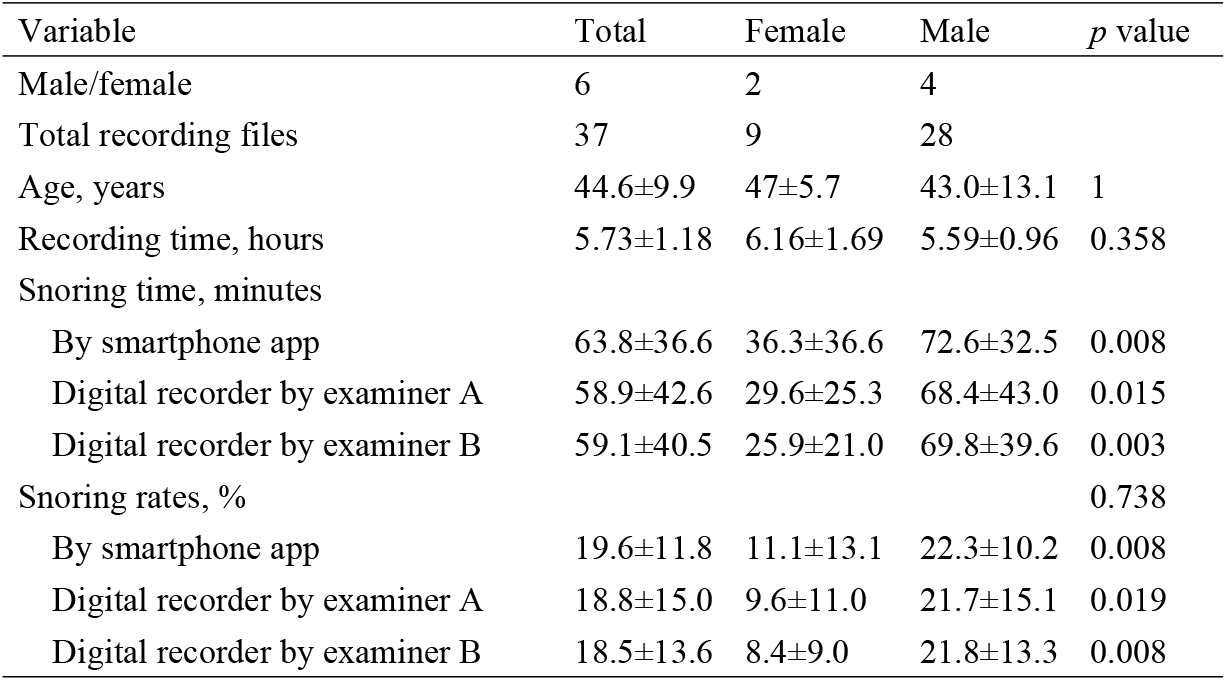
The demographics of subjects

**Figure 1.**
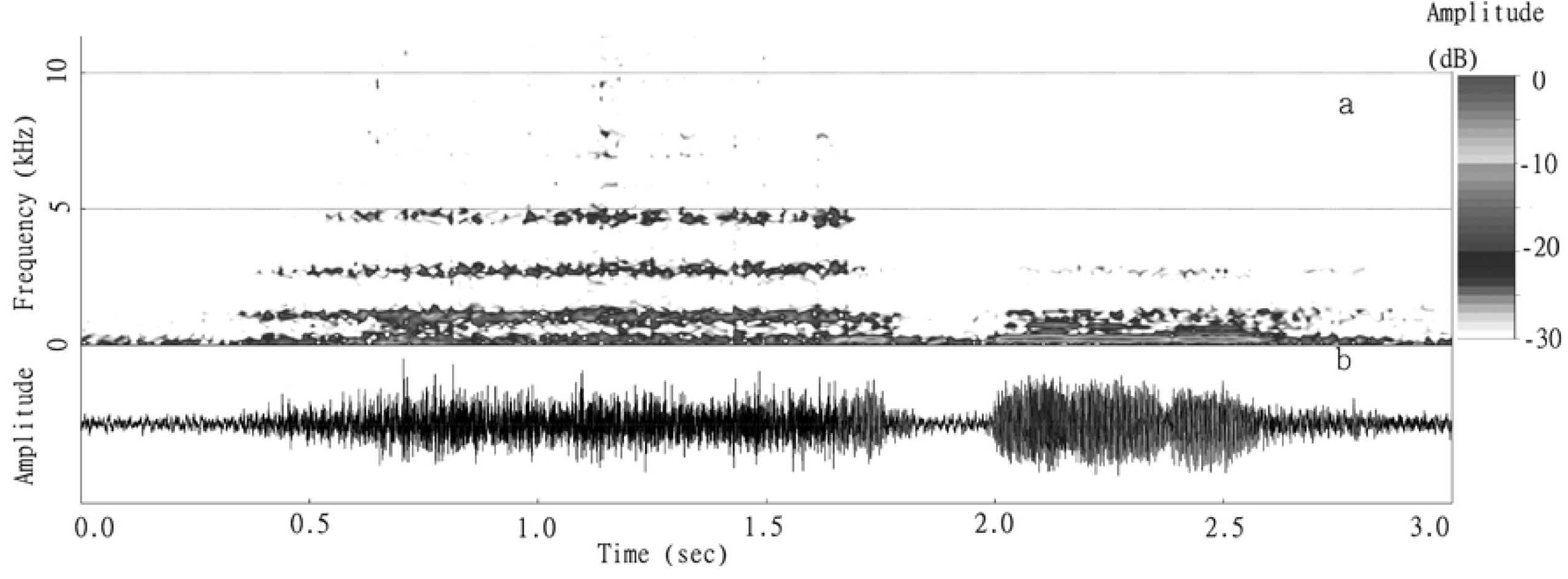
Example of a 3-s audio signal indicating snoring. The spectrogram of frequency (upper): three mean frequency bands of snoring (4694, 2713, and 1083 Hz) on the spectrogram; the fourth mean frequency band of snoring was hidden in background noise. The amplitude of snoring (button): the amplitude exhibited an ascension–descension pattern.

**Figure 2.**
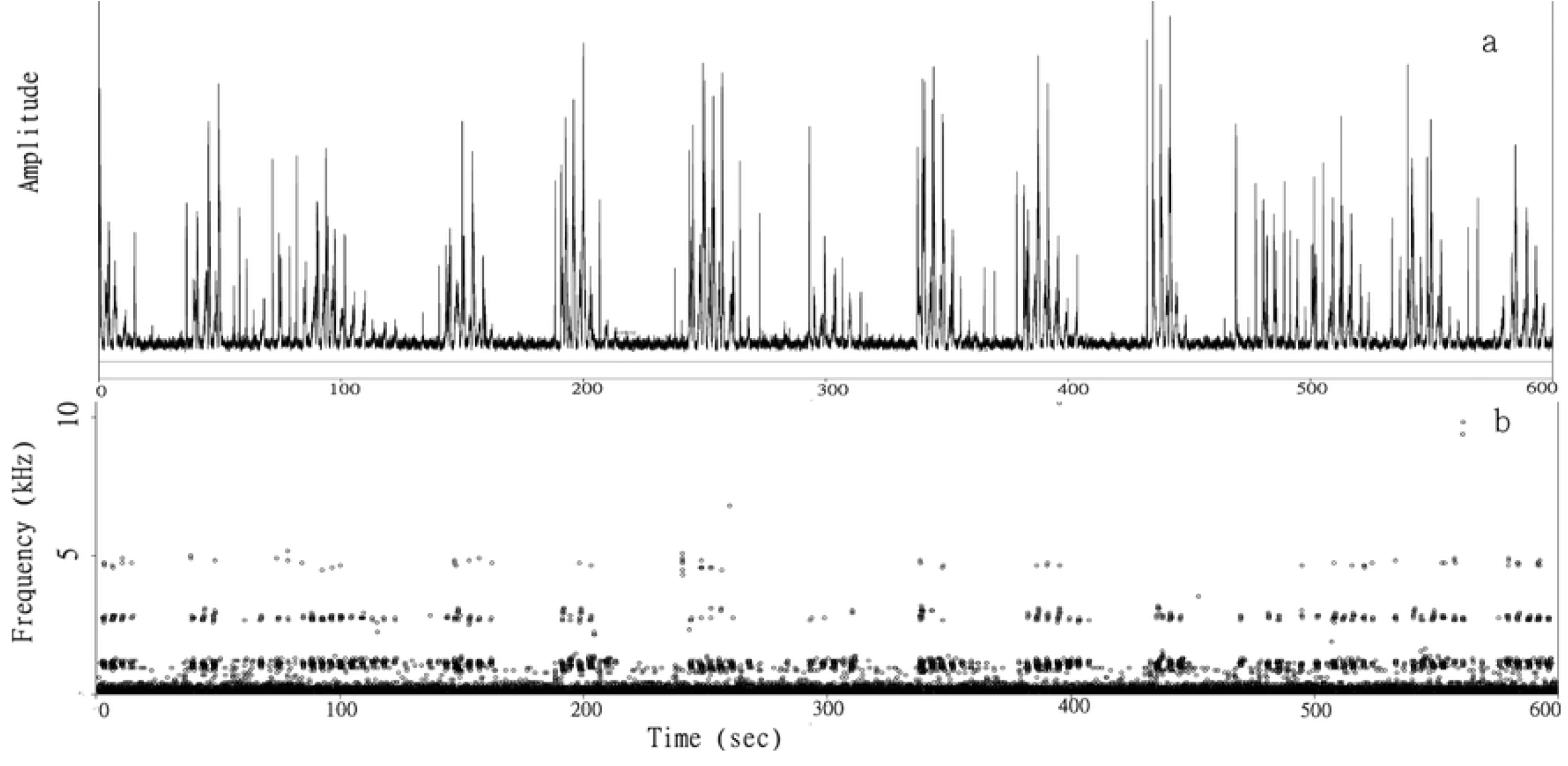
Example of snore interpretation by using the frequency domain and amplitude domain. A 600-s typical snoring duration including signals and intervals. The graph shows grouped near-isometric spikes with specific corresponding frequency bands. The clustered spikes exhibit a unique ascension–descension pattern regarding its amplitude, with intervals (typically less than 100 s) in between.

**Figure 3.**
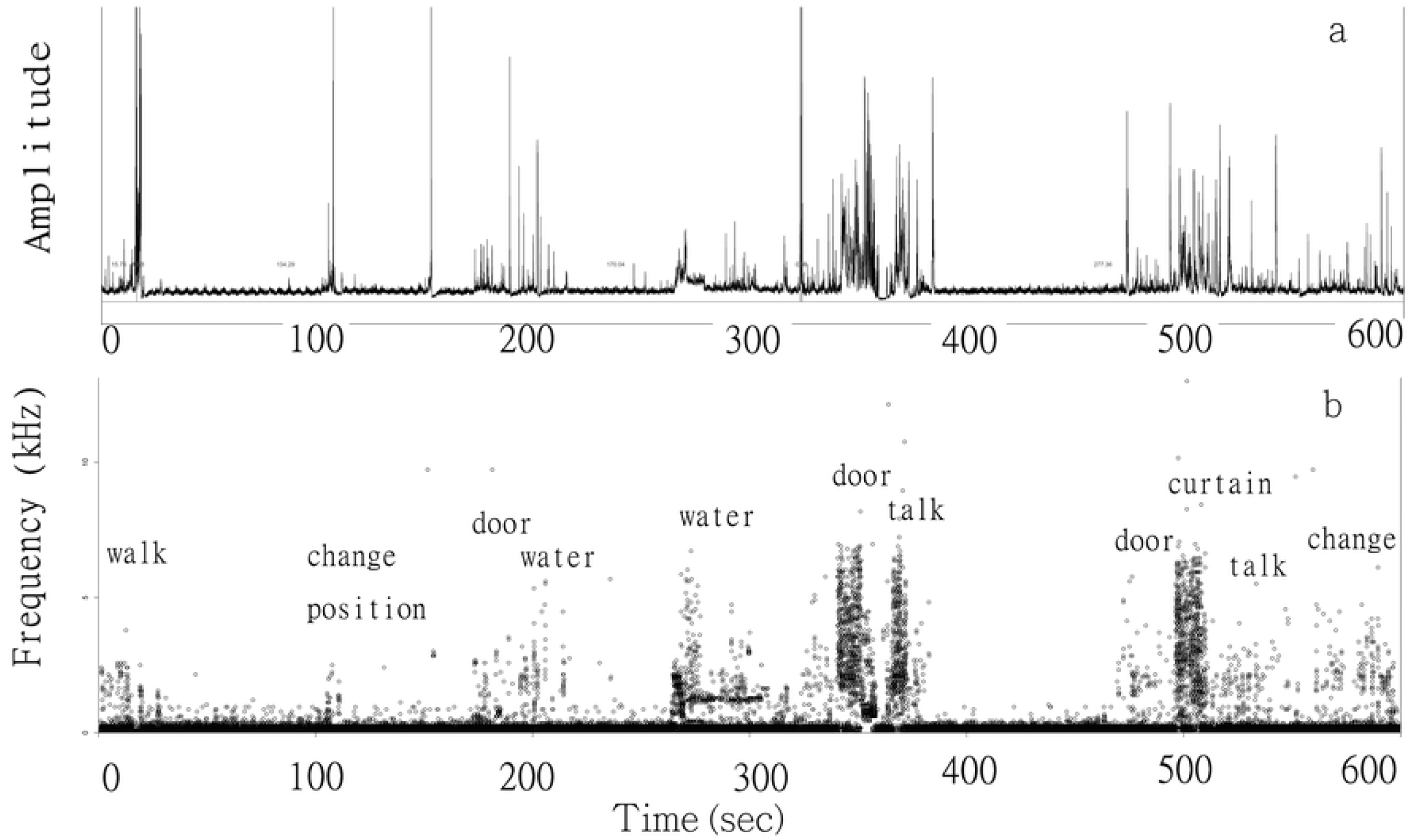
Examples of noises such as those from walking, changing position, doors opening or closing.

In this study, men had significantly greater snoring durations and rates than did women. To test for the reproducibility of our algorithm in calculating snoring durations, the authors examined the files digitally recorded by Examiners A and B. The differences in the scatterplots of snoring rates between those produced by the SnoreClock app and those obtained by Examiners A and B are shown in Figures 4a–c. The correlative coefficient of snoring rates calculated by Examiners A and B was 0.865 (*p* < 0.001), which is higher than that calculated by the SnoreClock app (Table 2). The difference between the means of snoring rates calculated by the two examiners was small. Additionally, both examiners obtained values that were lower than those obtained by the SnoreClock app. However, the three calculated snoring-rate values exhibited no statistically significant difference (*p* = 0.738; Figure 5).

**Table 2.**
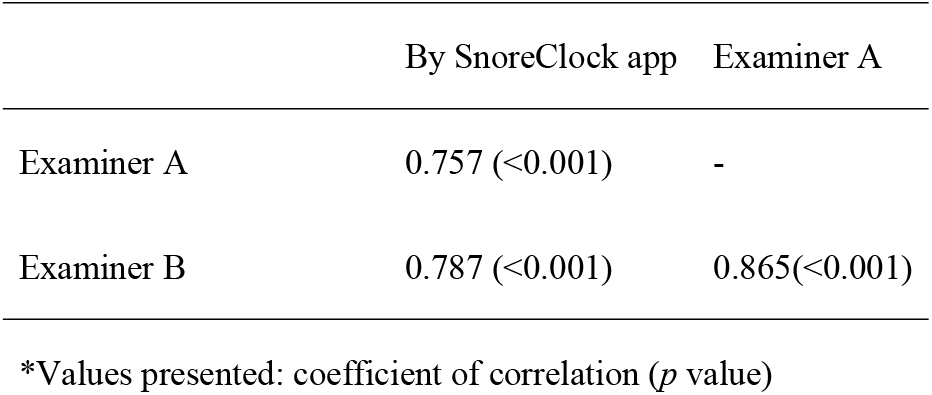
Correlation of snoring rates by different methods

**Figure 4.**
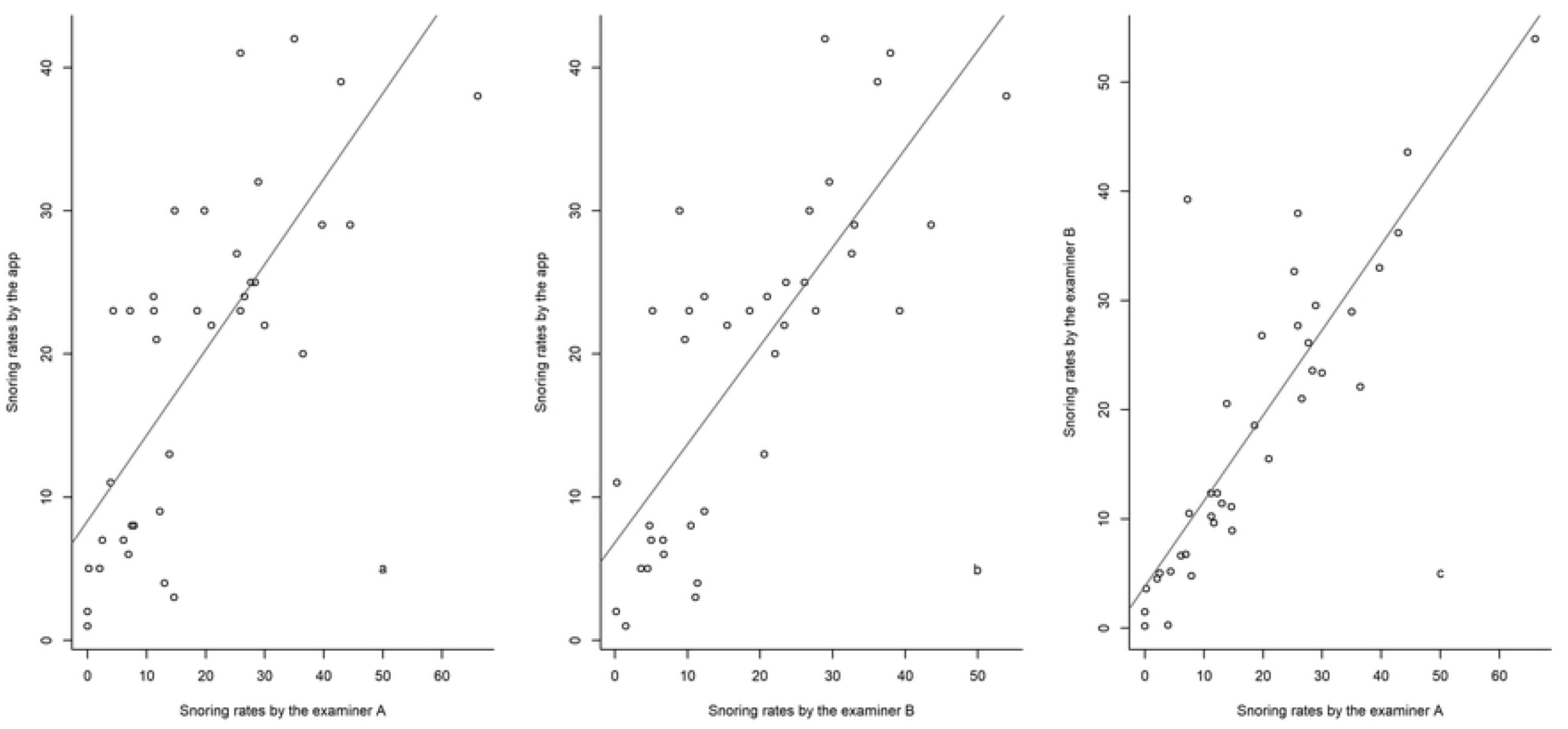
Scatterplots of snoring rates detected using the SnoreClock app and digital recorder files, as interpreted by Examiners A and B.

**Figure 5.**
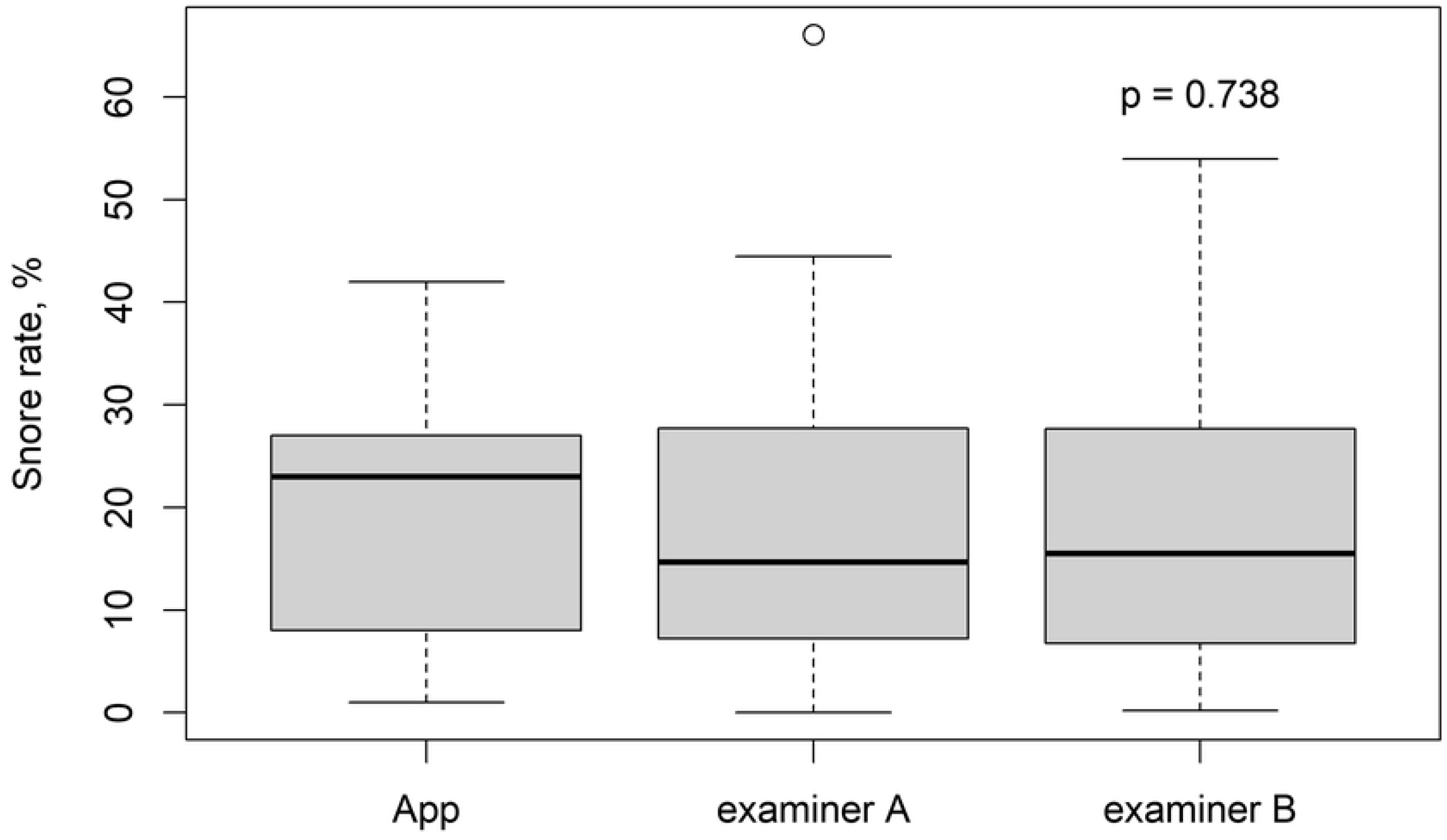
Boxplot showing snoring rates detected using the SnoreClock app and digital recording files, as interpreted by Examiners A and B.

## 4. Discussion

In the current study, snoring duration was determined according to signals and intervals (typically within 100 s). The featured signals of snores in the amplitude domains shared a similar ascension–descension pattern. Through Hilbert transformation, each individual ascension–descension snoring signal unit was processed to identify individual spikes, which were subsequently grouped to form near-isometric spike clusters. The intervals between clustered spikes were then determined. In the frequency domain, one or more fixed corresponding frequency bands in the snoring signals were also observed.

In this study, the authors could visualize snoring durations according to typical spectrograms from the audio files and thereby calculate the duration of snoring for each file. Noise patterns were easily identified and visualized because of their differences from the aforementioned typical features of snoring. With the use of a digital recorder at 44.1 kHz and R software, this dependable algorithm can be reliably used again. The correlation coefficient of snoring rates calculated by Examiners A and B was higher than that calculated by the SnoreClock app. The difference between the mean snoring rates calculated by the two examiners (A or B) and the SnoreClock app was small and statistically nonsignificant (*p* = 0.738). Therefore, people with habitual snoring can reliably use smartphone apps, such as SnoreClock, to determine rates of snoring duration at home.

For validation of snoring sounds, the acquired audio files were digitalized at a sampling frequency of 44.1 kHz, PCM, and 16 bits per sample, in accordance with previous research.[9] In Dafna’s study, all audio files were downsampled to 16 kHz, and each audio file underwent an adaptive noise suppression process based on the Weiner filter. Accordingly, they could detect snoring sounds with an accuracy of >98%.[9] However, the authors did not perform down sampling, and the noise durations presented were not suppressed. In our study, the authors validated snoring sounds on the spectrogram by identifying matching typical amplitude and frequency domains. Snoring signals generally exhibited one or more discrete frequency bands, and their spiked amplitudes occurred in groups at characteristic near-isometric intervals. Our dependable algorithm can be considered reliable for detecting snoring durations through visualization.

The analysis of snoring through audio recording with a microphone was applied and serves as a convenient option. Studies have reported the advantages of audio recording as its unobtrusiveness, portability, and low cost. [14, 22] However, data sets of audio recordings require further analysis to define snoring durations because this algorithm is inaccessible for people who did not have R software applications.

Smartphone apps provide convenient and personalized sleep care, and their accuracy in predicting snoring rates ranges from 93% to 96%.[18] Nevertheless, snoring does not have fixed and constant audio characteristics; the authors compared the rates of snoring durations between smartphone apps and sound recorders. Studies have reported that snoring duration measurements obtained using smartphone apps are highly similar with those obtained through polysomnography.[23] In the current study, the authors compared the spectrograms of audio files obtained using a smartphone app (SnoreClock) with those obtained using the proposed dependable algorithm for detecting snoring duration.

### 4.1 Limitation

This study has the following limitations. First, because of the small sample size of 36 files from six subjects, further study with a large sample size is warranted to confirm the results. Second, the time spent before falling asleep was not omitted; therefore, the rates of snoring durations may have been underestimated. Finally, snoring apps might be unable to distinguish between snoring sounds from different individuals in the same room.

## 5. Conclusion

The examiners determined a higher correlation than those between the SnoreClock app’s calculation and digitally recorded files (by a Sony recorder). Smartphone apps, such as SnoreClock, may provide a dependable algorithm that can be conveniently used at home to measure snoring rates. In the present study, the authors provide a truly dependable algorithm for detecting snoring sounds for researchers interested in investigating snoring.

## Author Contributions

**Conceptualization:** Hsueh-Hsin Kao, Yen-Chang Lin, Yee-Hsin Kao, Chun-Lung Wang, Jui-Kun Chiang

**Data curation:** Yen-Chang Lin, Madan Ho, Hsiao-Chen Yu, Jui-Kun Chiang

**Formal analysis:** Jui-Kun Chiang

**Funding acquisition:** Jui-Kun Chiang

**Investigation:** Hsueh-Hsin Kao, Yee-Hsin Kao

**Methodology:** Yee-Hsin Kao, Jui-Kun Chiang

**Project administration:** Yee-Hsin Kao, Jui-Kun Chiang

**Resources:** Jui-Kun Chiang

**Supervision:** Hsueh-Hsin Kao, Yee-Hsin Kao, Jui-Kun Chiang

**Validation:** Yee-Hsin Kao, Jui-Kun Chiang

**Visualization:** Hsueh-Hsin Kao, Jui-Kun Chiang

**Writing – original draft:** Hsueh-Hsin Kao, Yee-Hsin Kao, Jui-Kun Chiang

**Writing – review & editing:** Hsueh-Hsin Kao, Chun-Lung Wang, Yee-Hsin Kao, Jui-Kun Chiang. Hsueh-Hsin Kao and Yee-Hsin Kao contributed equally. Chun-Lung Wang and Jui-Kun Chiang were the co-respondent authors.

